# Age Classification of White-tailed Deer Via Computer Vision and Deep Learning

**DOI:** 10.1101/2025.07.01.662304

**Authors:** Aaron J. Pung

## Abstract

Accurate age estimation of wild whitetail deer remains a significant challenge for wildlife management. This study presents the first application of computer vision to whitetail buck age estimation using trail camera imagery, evaluating over sixty classification algorithms from traditional machine learning to advanced deep learning techniques. Our approach utilizes transfer learning and CNN ensembles to achieve a breakthrough cross-validation accuracy of 76.7 ± 5.9%, substantially outperforming traditional classifiers (57%), human expert assessment (60.6%), and morphometric methods (63%), and surpassing the 70% accuracy threshold required by professionals for wildlife management decisions. Furthermore, attention map analysis of the ResNet-18 ensemble reveals that the model learns to focus on the same anatomical features (neck, chest, and stomach regions) that human experts rely upon for age assessment. This biological validation demonstrates that the CNN identifies genuine age-related morphological changes rather than spurious correlations, lending credibility to its predictions. This breakthrough offers wildlife professionals a practical tool to dramatically reduce manual age assessment workload while exceeding current accuracy standards, potentially transforming how deer populations are monitored and managed across North America.

## 1 Introduction

Collecting and analyzing images of local fauna is crucial to the success of wildlife management and conservation efforts. State-of-the-art motion-activated cameras enable wildlife professionals and enthusiasts to photograph animals regardless of date, time, or weather. In turn, the images can be transferred to a computer for visualization, analysis, and insights into population health and behavior.

Of the many species commonly photographed in North America, whitetail bucks (*Odocoileus virginianus*) are particularly captivating due to their majestic morphology and changes in their appearance as they mature. Nonetheless, accurately predicting the age of a whitetail buck can prove difficult. To predict the age of living bucks, many hunters and enthusiasts turn to “aging on the hoof” (AOTH). Found in literature as early as 1978[1], this method attempts to predict the whitetail’s age based on the deer’s location, photograph date, and body traits such as the chest, stomach, neck, legs, and antlers.[2]– [6]

Previous research efforts have been performed on captive, known-age deer. Flinn’s 2010 study[6], for instance, analyzed 64 morphometric ratios of living, wild white-tailed deer across three states: Mississippi, Louisiana, and Texas. The study resulted in two age prediction models, one simple and one complex. When applied to individual age classes, the simple model achieved 33% accuracy for pre-breeding periods and 40% accuracy for post-breeding periods. The complex model, on the other hand, achieved 53% accuracy for pre-breeding periods, and 63% accuracy for post-breeding periods.

Another study in 2013 by Gee et al.[7] found that wildlife enthusiasts and professionals achieved a 36% accuracy rate when aging deer on the hoof, although individual scores widely varied, ranging from 16% to 56%. Furthermore, prediction accuracy was found to decline as the age of the buck increased. For instance, wildlife professionals performed better at predicting the age of yearling bucks (1.5 years) than older whitetail bucks (ex. 4.5 years). Despite the relatively low prediction accuracies, the same wildlife professionals believed a minimum accuracy of 70% is needed to be useful in wildlife management decisions, and a minimum accuracy of 80% is needed for research applications – standards that neither morphometric models nor human assessors have achieved.

The ages of whitetail bucks can be professionally estimated in many different ways. The first is by contributing trail camera images or video to established organizations like the National Deer Association (NDA), in which a panel of wildlife biologists and experienced professionals reach a mutual consensus over each photo (Lindsay Thomas, personal communication, May 1, 2025). To be considered, the deer in the contributed photograph must meet several requirements: it must be broadside, standing relatively still, have its head up, the image must be well lit and not blurry, it must have its entire body contained in the image frame, it must be photographed in pre-rut or during rut, and there can be no velvet on the buck’s antlers.

Even with these requirements in place, a large request volume remains, overwhelmimg wildlife experts and making it difficult to give equal attention to each submission. On the other hand, if the buck is deceased its jawbone and teeth can be removed and submitted to a laboratory for further age analysis[8]–[10]. Although more scientific than AOTH, this process can still prove cumbersome and is not always feasible depending on the state of the deer’s body.

To alleviate the workload on wildlife professionals and enhance awareness within the whitetail community, many organizations provide tutorials on best AOTH practices. In each tutorial, highly trained professionals discuss general body characteristics and aging techniques across different age classes, often providing clear examples to help clarify the message. Over time, the growing database of information, tutorials, and images helps viewers develop an intuition for aging whitetails that can be applied in their own experiences.

Much in the same way that humans learn to predict age based on a deer’s body shape, Machine Learning (ML) and Computer Vision (CV) can be leveraged to generate age prediction models based on the same database of trail camera imagery and pre-determined ages. Models like Convolutional Neural Networks (CNN) automatically detect and extract features within trail camera images, removing the manual labor of feature identification exemplified in Flinn’s 2010 thesis. This study reports the first known use of computer vision and CNNs in predicting the age of whitetail bucks from trail camera imagery, offering wildlife experts, hunters, and enthusiasts a new tool in wildlife management and conservation efforts.

## 2 Human Vision

### 2.1 Data

As a part of its ongoing mission to engage and educate outdoor enthusiasts, the NDA publishes different forms of media discussing whitetail age prediction. In their weekly online newsletter, an “Age This!” column presents readers with an image of a whitetail buck and a survey inviting the reader to estimate the age of the deer without seeing the “correct answer” (the NDA’s age determination) or the response of other readers. In addition to the image, readers are also given the date and location the image was taken, allowing them to correlate the deer’s size and shape with the local rut season.

Results of the survey are provided in the subsequent week’s newsletter along with the NDA’s age determination and a brief discussion of factors used to finalize the age prediction. Results from the “Age This!” survey were collected and analyzed over a six-month period to gauge the accuracy of outdoor enthusiasts in predicting the age of whitetail bucks via AOTH.

### 2.2 Prediction Accuracy

Analysis plots of the survey data are shown in Figure 1. The bar chart on the left shows the reader’s prediction accuracy by age class; white percentage values on each bar indicate the percentage of enthusiasts who correctly guessed the deer’s age. For example, on average over six months of data, 79.7% of readers correctly guessed the age of yearling bucks, while 60.7% of readers correctly identified the age of 2.5 year old deer.

**Figure 1.**
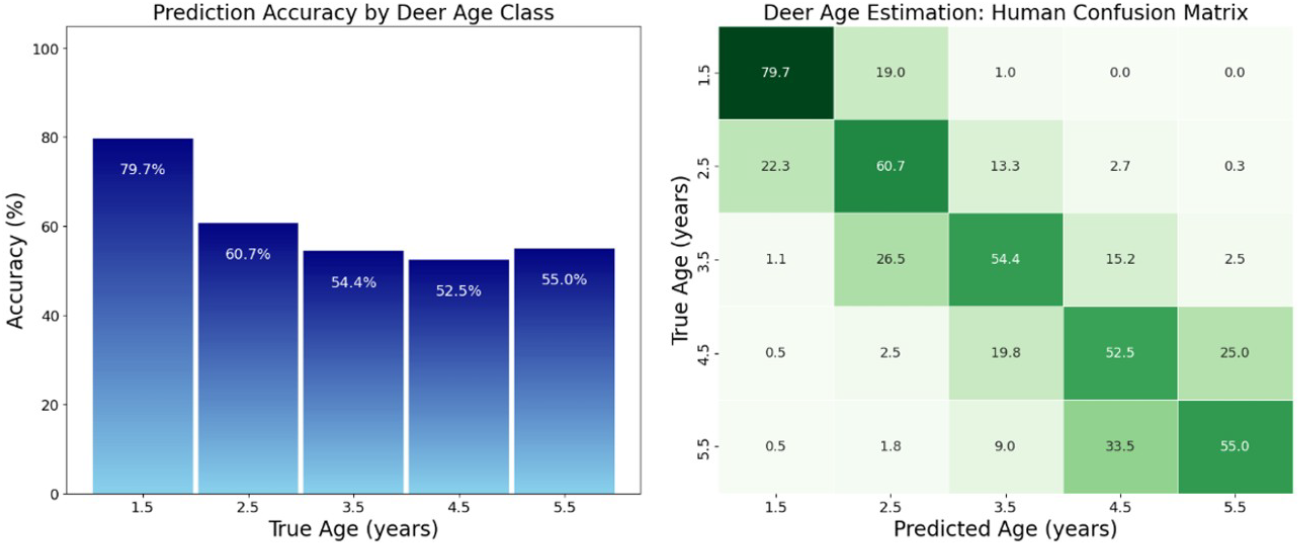
Human prediction accuracy for each age class are presented in the form of (left) a bar chart, and (right) a confusion matrix.

For each age class, the accuracy of readers’ estimates consistently lies below 80%, the metric professionals agree is needed for research purposes. For deer aged 2.5 years or older, accuracy rates fell well below 70%, the minimum metric needed for wildlife management purposes. The lowest accuracy predictions occurred with deer aged 4.5 years old (52.5%); this may be due to the buck’s maturing body shape, which changes significantly with location, nutrition, or time of year (ex. rut). Estimation accuracy tends to increase again for bucks 5.5 years or older. Since the final age class encompasses multiple ages (*≥* 5.5 years), however, readers may simply find it easier to place older bucks in this category without correctly predicting the age.

The table on the right contains the same information, but also characterizes readers’ incorrect guesses. Known as a “confusion matrix”, each row in the table represent one True Age value and its columns represent predicted values. For instance, a yearling buck (True Age of 1.5 years) is correctly predicted to be 1.5 years old 79.7% of the time – this is indicated in the matrix as the dark green square in the upper left corner. However, the same yearling buck is incorrectly predicted to be 2.5 years old 19.0% of the time, represented by the light green box in the top row, second from the left.

Further analysis of the human confusion matrix reveals intriguing insights into trends in human AOTH predictions. For instance, bucks with a true age of 2.5 years are predicted to have the correct age 60.7% of the time – but when the guess is incorrect, the prediction is twice as likely to represent a younger buck. The same is true for 3.5 year old bucks, but not true for 4.5 year old bucks; in this case, inaccurate predictions tend to estimate an older age.

### 2.3 Analytics

Individual prediction accuracy (*IPA*) can be calculated by averaging the percentage of correct age estimates (*P*_*i*,*correct*_) over all images[11] using equation (2.1).

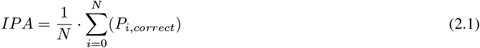

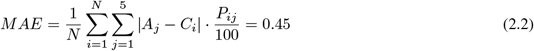

The average error across all images can be found via the mean absolute error (MAE), defined by equation (2.2). Here, *A*_*j*_ represents the *j*^*th*^ age class (1.5, 2.5, etc.), *C*_*i*_ represents the correct age class for image *i*, and *P*_*ij*_ represents the percentage of individuals who estimated age class *A*_*j*_ for image *i*. Strictly speaking, the MAE is the weighted average of absolute errors across all age classes for each image.[12], [13]

The values of *IPA* (0.606) and *MAE* (0.45) indicate that the average enthusiast is correct 60.6% of the time and their incorrect guess differs from the NDA’s age estimate by less than half a year. It should be noted, however, that even NDA panelists have been known to change their estimates over time for the exact same image.

Although the “Age This!” survey collects individual age votes, groups of voting individuals would be able to discuss ideas and image details, arriving at a different conclusion than what an individual might reach on their own. Similar to individual accuracy, majority accuracy is found by calculating the percentage of time the majority vote is correct. Of the available data samples, the majority’s vote was incorrect five times, resulting in a majority accuracy of 79.3%, an improvement of 18.7% over the average individual.

## 3 Dataset and Methodology

### 3.1 Data Collection

The image database used in this study contains 457 images of whitetail bucks collected from 13 independent sources, each containing one or more highly experienced wildlife professionals. To ensure consistency with professional age estimation standards, only photographs from the NDA with verified age assessments were included. Within this subset, only colored images were retained for analysis given the change of fur color as deer age.

Although the standardization process reduces the dataset to 197 images, it also ensures that all training data meets the quality standards used by wildlife professionals for age determination. The size of the dataset also reflects the realities of wildlife research where obtaining large quantities of professionally-verified data is challenging and expensive.

Each image undergoes a standardized preparation process (Figure 2). The original photograph is cropped to a square format that captures as much of the deer as possible. The cropped images are resized and subsampled to fit the expected model input size. This standardization ensures the CV model locates and evaluates each deer based solely on their physical characteristics like body proportions, antler development, facial features, and muscle definition rather than being influenced by differences in photo quality, lighting, or background elements. A subset of images has been included in Figure 3.

**Figure 2.**
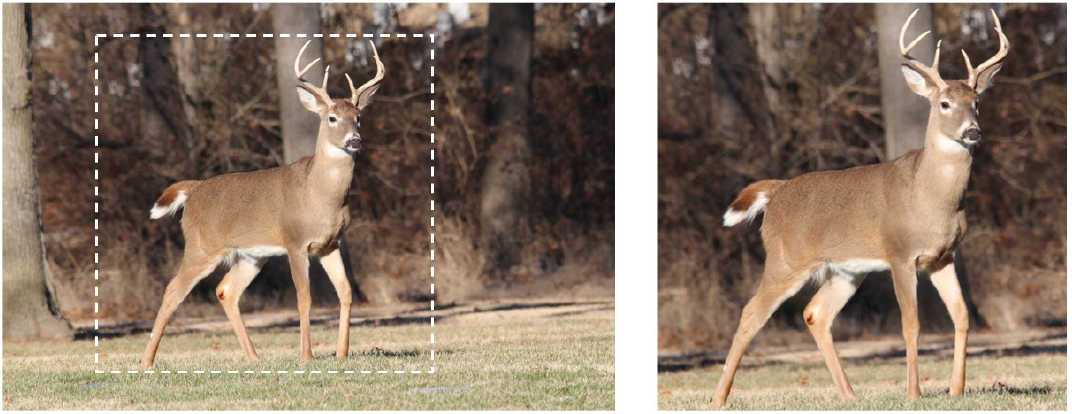
(Left) A trail camera image of a whitetail buck. The dashed white box represents the boundary of the crop used to achieve (right) the standardized square image.

**Figure 3.**
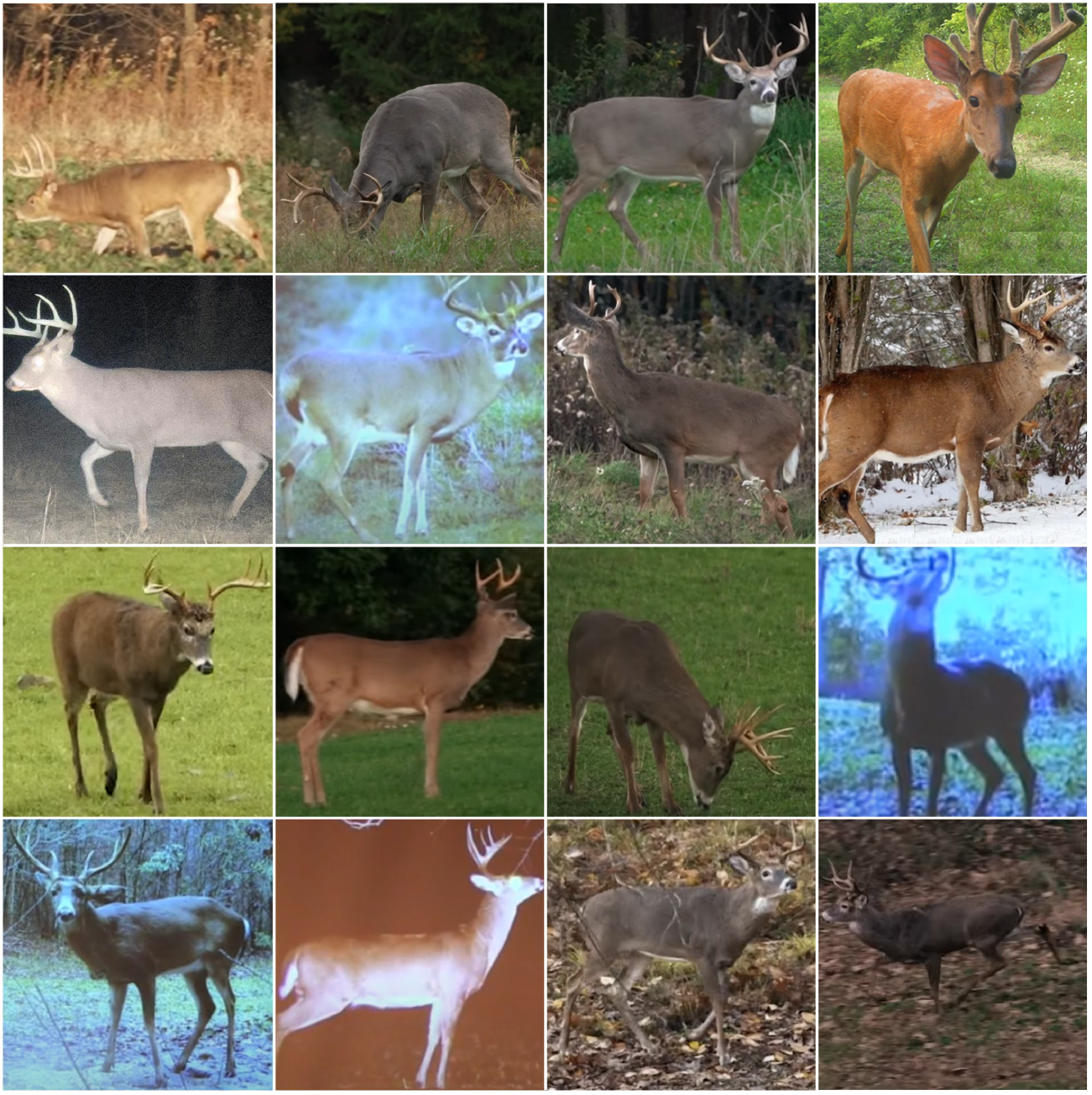
A sample of images from the standardized dataset.

In addition to the survey, images were collected from videos, online posters, and other media produced by the NDA, resulting in a dataset with wide geographic variation (¿14 states) and less stringent requirements than the survey itself.

Deer in the database compiled for this study need not be broadside, and do not need to have their head up. Images can be blurry, lighting can be poor, and a portion of the deer’s body may be left out of the image. Image metadata like date and location are never fed into the model. Furthermore, none of the images contain artificial borders or digital artifacts, so any classification accuracy achieved by the model reflects its ability to recognize genuine biological age markers that hunters and wildlife biologists use in practice.

### 3.2 Issues and availability

Following image standardization, two age-related issues remain, the first of which is unequal age representation. Since fawns and yearlings are comparatively easier to age, their images do not require as much scrutiny as older deer and their images do not appear as often on institution websites, tutorials, or other similar media. Similarly, community-wide interest tends to focus on mature bucks that are old enough to harvest. Coincidentally, these older bucks are also more difficult to age, so their images gain more attention with hunters, enthusiasts, and professionals alike. Both factors heavily skew the age distribution towards mature bucks.

The second issue is a lack of precision in how estimated ages are recorded. The age of a whitetail buck is often estimated in half-year increments (ex. 1.5, 2.5, etc.) due to the deer’s life cycle; newborn fawns are born in the Spring and mature bucks are often harvested during the Fall hunting season. Between 1.5 years and 5.5 years, the buck’s body changes and grows in predictable patterns. Even though the buck’s body continues to change after 5.5 years, changes may not be as evident; for this reason, bucks older than 5.5 years are typically grouped together under the heading of “5.5+ years”. Accordingly, bucks within the dataset used for this study are artificially labeled as “5.5 years” even if their professionally determined age exceeds 5.5.

### 3.3 Age and distribution

The dataset used in this study comprises 197 colored images and an equal number estimated age values. Age class distribution is as follows: (1.5 years) 30 images, (2.5 years) 36 images, (3.5 years) 36 images, (4.5 years) 52 images, and (5.5+ years) 43 images. Twenty percent of the data (40 images) are set aside as a “test” dataset and the remaining 80% (157 images) form the “training” dataset. The test set maintains proportional age class distribution with 6 samples of 1.5-year bucks, 7 samples each of 2.5 and 3.5-year bucks, 11 samples of 4.5-year bucks, and 9 samples of 5.5+ year bucks, ensuring representative evaluation across all age categories.

### 3.4 Augmentation

Like humans, machine learning models perform better with larger datasets – the more images a model can learn from, the more accurate it can become. Classification problems like this one typically require at least 1,000 samples per category, more than 25 times the samples collected for this study. To reach the ideal number of images, datasets can be artificially expanded through augmentation.

By applying subtle modifications to the training data, more images can be obtained without altering the deer’s characteristics or body proportions. But it is important that modifications made during the augmentation process mirror natural variations hunters and wildlife observers might encounter in the field. For instance, adjusting brightness simulates changes in sunlight throughout the day, while cropping an image mimics a deer walking out of the view of the camera. Small rotations can be used to mimic imperfect camera mounting, and horizontally flipping an image mimics the same deer walking in the opposite direction along a trail. Since image augmentation can applied independently across all age classes, larger datasets can be obtained with perfectly balanced age classes.

However, over-augmenting can destroy the very features that distinguish each age class: excessive cropping might eliminate key body characteristics, while extreme brightness changes could make body details or facial features unrecognizable. The goal with augmentation is to enhance the dataset while preserving the biological markers that experienced hunters and biologists rely on for age estimation. In this study, augmentation occurs prior to the training process, and is used equally across all models. In total, applying rotations, horizontal flipping, varying lighting and contrast, and adding noise increased the training set by a factor of 40 such that each class of training data consisted of 1,680 images.

## 4 Computer Vision

### 4.1 Traditional Classification Models

During model analysis, twenty traditional machine learning algorithms were evaluated, including ensemble methods (Random Forest, AdaBoost, Extra Trees, Gradient Boosting), tree-based methods (Decision Trees, XGBoost), distance-based methods (K-Nearest Neighbors, Support Vector Machines), and linear models (Logistic Regression, Linear Discriminant Analysis).

The highest-performing traditional classifier was Random Forest, achieving 57.2% accuracy on the validation set. However, all traditional approaches are ultimately limited by the need for manual feature engineering and struggled to capture the complex visual patterns that distinguish age classes in trail camera imagery. This performance ceiling led to exploration of deep learning approaches that could automatically learn relevant features from the raw image data.

### 4.2 Transfer Learning via CNNs

In contrast, transfer learning is an ML technique that leverages knowledge from one domain to solve problems in another. For image-based applications, large Convolutional Neural Networks (CNNs) like EfficientNet[14], DenseNet[15], and ResNet[16] offer pre-trained parameters based on the ImageNet database.

ResNet-18 was selected as the primary architecture based on preliminary performance assessments, demonstrating a crossvalidation accuracy of 66.0% ± 7.2%. Although subsequent evaluation revealed ResNet-50’s superior cross-validation performance (76.7 ± 5.9%), ResNet-18 remained the focus of detailed analysis due to its faster training and lower computational cost.

The 4-layered ResNet-18 model is illustrated in Figure 4. The convolution layer (Conv1) and batch normalization (BN) are frozen, maintaining the CNN’s initial feature extraction. Layers 1, 2, and 3 are also frozen, preserving low-level edge and texture features, mid-level pattern features, and higher-level shape features, respectively. Allowing the fourth layer to be trainable enables the CNN to train exclusively on the augmented standardized data, extracting high-level semantic features such as relationships between the deer and its surrounding scenery. The fully connected layer (FC) is also trainable, ensuring the model output is optimized with the five pre-determined age classifications instead of the initial 1,000 image classifications used to originally train the ResNet model.

**Figure 4.**
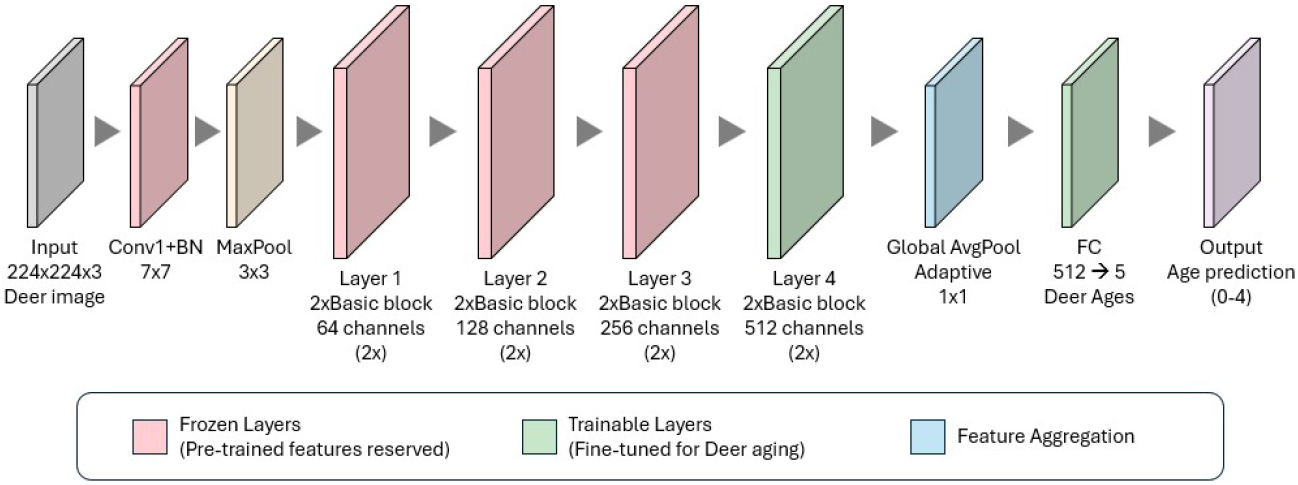
Architecture of the ResNet-18 model

### 4.3 Training and Configuration

The ResNet-18 training process is illustrated in Figure 5. Following separation of the training and test data, the test data is isolated and the training data is further separated into five stratified folds, ensuring proportional representation of each age class across all folds. Each fold is further split into a training dataset (125 images) and a validation dataset (32 images). The training dataset is then augmented to produce a total of 5,000 images.

**Figure 5.**
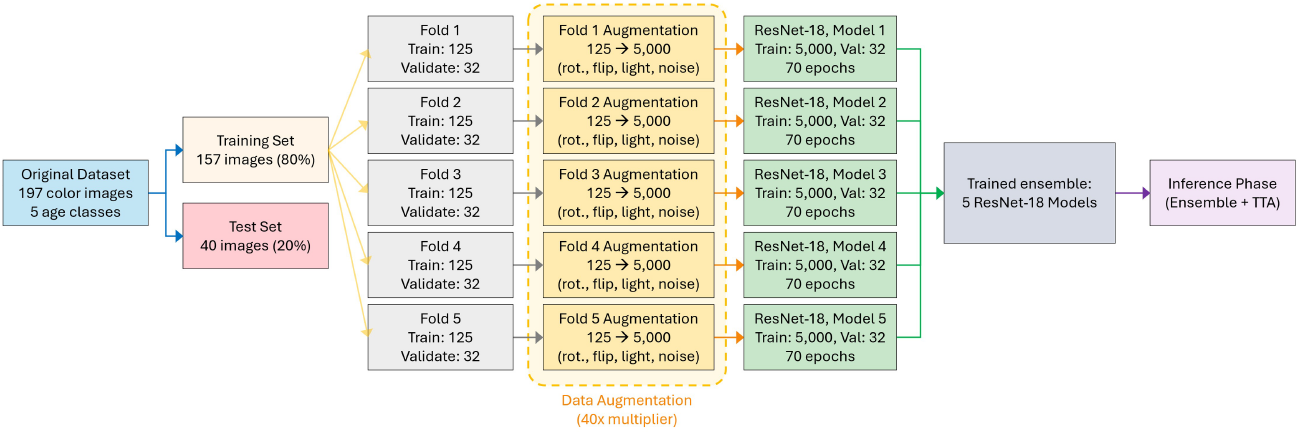
The process for training the ResNet-18 ensemble

Each of the five ResNet-18 models were trained using AdamW optimization with differential learning rates, 0.0003 for frozen backbone layers and 0.001 for the trainable classifier head. Learning rates followed exponential decay (*γ* = 0.95) with early stopping implemented to save time, using a patience of 20 iterations (epochs). Label smoothing (*α* = 0.1) provided additional regularization to prevent overfitting on the limited dataset. Cross-entropy loss was used as the primary objective function. Training typically converged after approximately 40 epochs per fold, with total training time of approximately 45 minutes on NVIDIA RTX 2060 hardware.

The augmented training data in each fold is used to train a separate ResNet-18 model. Each model trains independently on its fold’s augmented data and validates on the corresponding validation dataset over 70 epochs. After training is complete, the five trained models are combined into an ensemble. Once their weights are optimized, these models are used in the inference phase where Test-Time Augmentation (TTA) is applied. TTA is a technique in which each model makes predictions on both the original input image and a horizontally flipped version, then averages these predictions to improve robustness and accuracy.

### 4.4 Inference

Illustrated in Figure 6, the inference phase classifies each of the original (non-augmented) test images. Each input image and its horizontally flipped version are passed to all five fold-specific ResNet-18 models simultaneously. Within each model, TTA averages the probability distributions from the original and flipped images, and the resulting five TTA-averaged predictions are then ensemble-averaged using softmax normalization to produce a final probability distribution over age classes. The predicted age class is determined by selecting the highest probability, and these predictions are tabulated to calculate accuracy metrics for the entire ensemble model.

**Figure 6.**
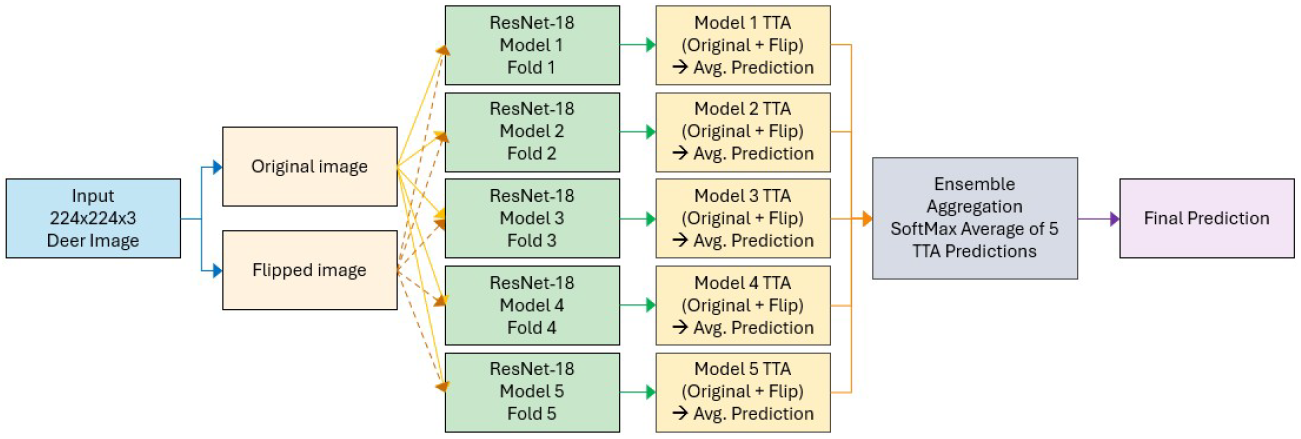
Inference process for the ResNet-18 ensemble

### 4.5 Results

Somewhat similar to the accuracy metrics presented in Figure 1, ML model accuracy is characterized by precision, recall, and F1-score. Precision measures the reliability of the model’s predictions for a specific age class; for example, of all the deer the model predicted as a particular age (e.g., 3.5 years), precision indicates the percentage that actually were 3.5 years old. Recall measures the model’s ability to find all deer of a specific age class; for example, of all the deer that actually were a particular age, recall characterizes the percentage the model correctly identified. High recall means the model successfully identifies most deer of the correct age group instead of misclassifying them as other ages. The F1-score provides a harmonic mean between precision and recall, offering a single metric that balances both the model’s reliability (precision) and completeness (recall) for each age class.

The precision, recall, and F1-score of the ensemble model for each age class are plotted in Figure 7. Across all age classes, predictions made by the ResNet-18 ensemble consistently exceeded 80%, well above the threshold determined by experts for use in management decisions as well as research applications. Even the more conservative cross-validation metric for both CNN models shows an improvement over previously reported models as well as human AOTH predictions.

**Figure 7.**
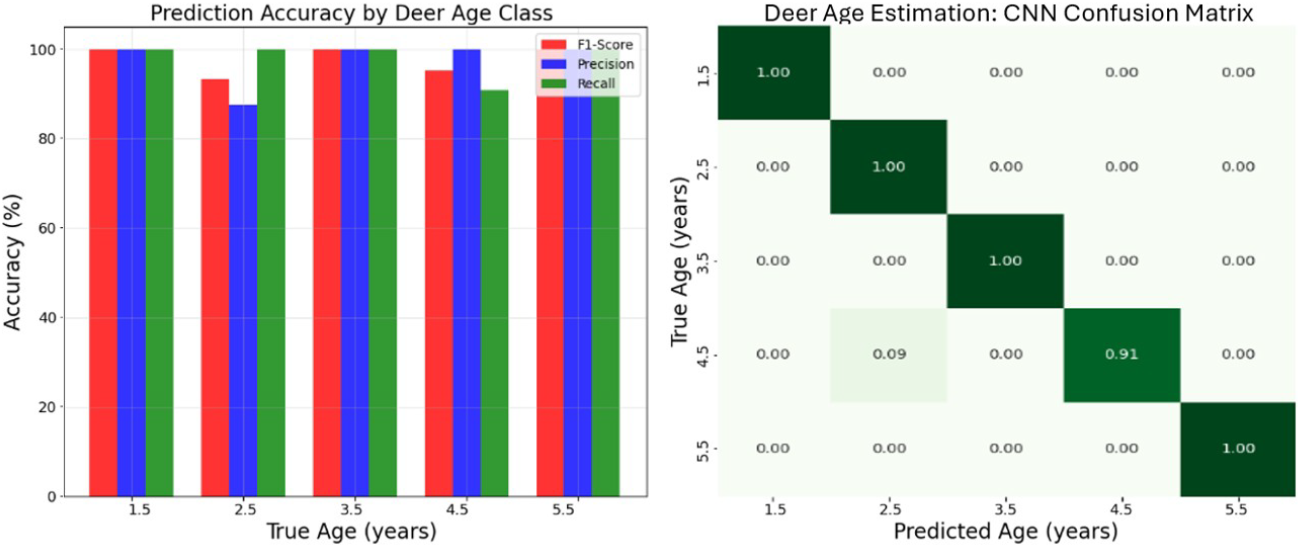
Accuracy metrics for the ResNet-18 ensemble are illustrated via (Left) F1-score, precision, and recall as well as (right) a confusion matrix.

The confusion matrix in Figure 7 illustrates the ResNet-18 ensemble achieving near-perfect classification, with only one misclassification: one 4.5-year buck incorrectly predicted as 2.5 years old. Interestingly, the ensemble’s misclassification occured with the same age group (4.5 years) that gives outdoor enthusiasts the most trouble. Perhaps more interesting is that when the ensemble predicted the incorrect age, the age it predicted (2.5 years) is one human enthusiasts were very unlikely to guess. Unlike outdoor enthusiasts, though, the model is fed neither date nor location information, so it is unable to correlate a given image with natural phenomena like rut.

Of equal significance is identifying features within each image the ResNet-18 ensemble uses to predict the buck’s age. Figure 8 illustrates the attention map for five cases, one for each age class. The true age of the deer (as determined by the NDA) is shown in the first column and the corresponding image of the deer is in the second column. The third column shows a low-resolution “attention map” highlighting regions of the image the CNN deems most important for a particular image. The fourth column overlays a high-resolution attention map with the original image.

**Figure 8.**
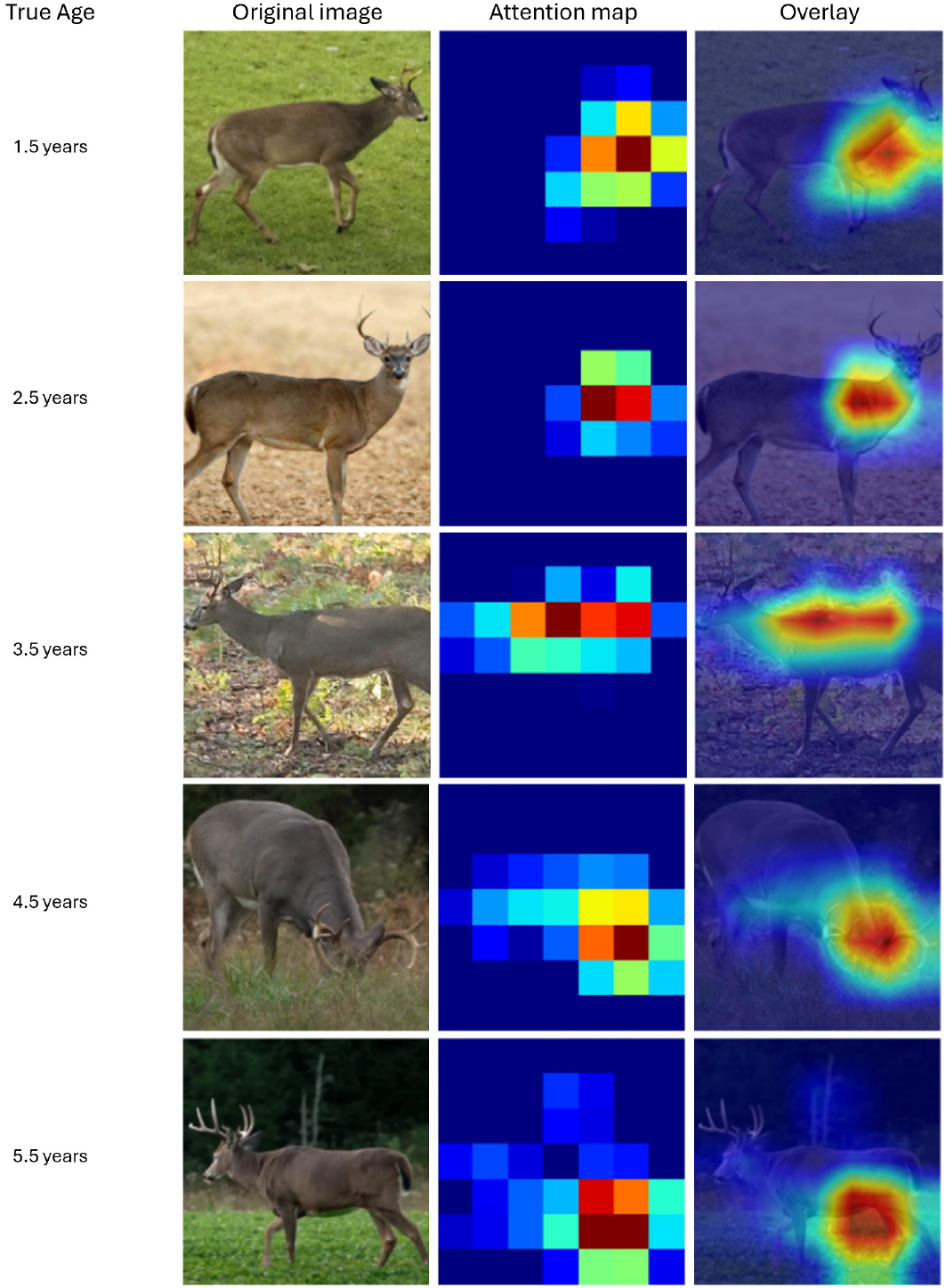
Attention map analysis showing (left to right): true age, original image, attention map, and overlay visualization for each age class.

Despite changes in the background clutter and the deer’s position, size, age, and posture, the attention maps for each image change in location, size, and shape demonstrating the model’s ability to pick up on purely biological traits within each image. Furthermore, the attention maps pick up on many of the same features that enthusiasts and hunters discuss when using AOTH. For instance, the attention map for the yearling buck in the top row seems heavily concentrated on the neck and chest – one of the more telling signs of a young deer. In contrast, the attention map for the 5.5 year old buck is heavily centered on the deer’s stomach – one of the main characteristics whitetail deer enthusiasts focus on when identifying older bucks. Furthermore, none of the attention maps for the five deer in Figure 8 focus on the deer’s antlers, aligning with another suggestion often cited by outdoor professionals.

## 5 Conclusions

This study explored the use of Computer Vision models in predicting the age of male whitetail deer based on trail camera imagery. Transfer learning with CNNs achieved mean cross-validation accuracies of 76.7%, representing significant improvements over traditional morphometric methods and human AOTH performance. While test set accuracies reached 97.5%, the more conservative cross-validation results provide a robust estimate for real-world deployment and exceed the 70% threshold considered useful for wildlife management decisions.

The model’s performance should be interpreted within the context of the limited dataset size inherent to wildlife research, and future work should validate these approaches across different geographic regions and camera systems.

